# MicroRNAome of *Spodoptera frugiperda* in Response to SfMNPV Infection

**DOI:** 10.64898/2026.08.12.744166

**Authors:** SM Gómez Bergna, LC Amorós Morales, A González Abad, J Vilches, SE Tongiani, R Salvador, V Romanowski, ML Pidre, ML Ferrelli

## Abstract

*Spodoptera frugiperda* is one of the most important agronomical pests due to its migratory capacity and broad host range. Since it is resistant to several insecticides, novel control strategies are being explored to control it. In this way, Spodoptera frugiperda Multiple Nucleopolyhedrovirus, a natural pathogen, has been proposed for its biocontrol.

In this work, we performed a small RNA-seq on uninfected larvae and larvae infected with SfMNPV to identify expressed miRNA, characterize them, and identify differentially expressed (DE) miRNA in the infected condition.

We identified several known and putative novel miRNAs, some of which are encoded in multiple copies and may be expressed within miRNA clusters. We also found 13 DE miRNA, most of them previously reported, two of them are putative novel miRNAs identified in this work. We predicted miRNA targets and found that their putative biological role could be related with processes relevant to the infection such as proliferative and apoptotic pathways, cell cycle regulation, autophagy, DNA damage response (DDR), vesicle transport, cytoskeleton remodelling, JAK/STAT and Toll signaling pathway, and immune response activation, among others. Moreover, we observed that several of the putative targets were hub genes in a predicted protein - protein interaction network. Finally, we found DE miRNA putatively associated with the regulation of viral gene expression, suggesting they might have a role in modulating the infection.

Our results contribute to better understanding the miRNA landscape in *S. frugiperda*, and their putative role upon SfMNPV infection.

## Introduction

*Spodoptera frugiperda*, commonly known as the fall armyworm (J. E. Smith, 1797)^1^ (Lepidoptera: Noctuidae), is one of the most significant agricultural pests worldwide due to its high migratory capacity and broad host range, capable of infesting over 350 plant species.^2^ Originally native to the Americas, *S. frugiperda* has now spread to every continent, prompting the Food and Agriculture Organization (FAO) to initiate a Global Action Plan to manage its impact.^3^ Current control strategies include chemical insecticides, transgenic crops utilizing *Bacillus thuringiensis* (Bt) technology,^4^ and cultural practices.^5^ However, resistance to both chemical insecticides and Bt crops has been increasingly reported.^5^

In response, alternative control strategies are being explored, including the use of entomopathogenic viruses, fungi, and other biological agents.^6^ Among these, Spodoptera frugiperda multiple nucleopolyhedrovirus (SfMNPV; species name: *Alphabaculovirus spofrugiperdae*), a member of the Alphabaculovirus genus (family *Baculoviridae*), has been proposed as an efficient biopesticide due to its capacity to naturally infect *S. frugiperda*.^7^ SfMNPV possesses a double-stranded DNA genome of approximately 140 kbp, encoding around 145 coding sequences, depending on the viral strain. Like other baculoviruses, SfMNPV has been extensively studied for its biological control properties,^8^ and has been approved for commercial use in several countries.^7^ Viral infection in insects involves complex interactions between the virus and host signaling pathways. Key antiviral responses include activation of the JAK/STAT, Toll/Imd, NF-κB, apoptosis, miRNA, and siRNA pathways.^9,10^ Additionally, signaling cascades such as ERK and PI3K/Akt may have context-dependent pro- or antiviral roles. Hormonal regulation, particularly through the 20-hydroxyecdysone (20E)/juvenile hormone (JH) axis, also influences infection outcomes.^11,12^

MicroRNAs (miRNAs) are small non-coding RNAs ranging from 19 to 24 nucleotides, which regulate gene expression by base-pairing with target mRNAs, triggering their degradation or translational repression.^13,14^ A single miRNA can target multiple mRNAs, enabling the fine-tuning of complex cellular processes.^15^ In insects, miRNAs are known to regulate diverse physiological processes, including development, oogenesis, metamorphosis, and immunity.^16,17^ Several miRNAs from *S. frugiperda* have been identified under various experimental conditions,^18–22^ and some have been implicated in responses to baculovirus infections in *S. frugiperda.*^23–25^ Similar findings were reported in studies on other insect species.^26–36^

However, to date, no studies have characterized the microRNAome of *S. frugiperda* specifically in response to SfMNPV infection. In this work we aimed to identify and bioinformatically characterize *S. frugiperda* miRNAs expressed in these conditions. We identified novel miRNAs and duplication events, predicted novel miRNA clusters, and characterized differentially expressed miRNAs and their putative biological role in *S. frugiperda* following the infection with an Argentinian isolate of the virus, SfMNPV ARG-M.^37^

## Materials and Methods

### *S. frugiperda* larvae infection with SfMNPV

*S. frugiperda* larvae were reared individually at 27°C, with 16 h light:8 h dark, and fed with artificial diet at the Instituto de Investigación Microbiología y Zoología Agrícola (IMyZA, INTA Hurlingham, Buenos Aires, Argentina). SfMNPV OB suspension was prepared as described by Masson et al (2021).^37^ Third-instar larvae were infected *per os* with 5×10^8^ OB/ml suspension of SfMNPV-M Argentinian isolate using the droplet method.^38^

### RNA samples preparation and sequencing

Total RNA was extracted from two infected (48 hpi) and two control, uninfected samples using Tri-Reagent (MRC), according to the manufacturer’s instructions. Each sample consisted of pools of ten larvae. RNA quantity and quality were verified using a NanoDrop 2000 spectrophotometer (Thermo Scientific). Also, RNA integrity was assessed on a 1 % (w/v) agarose gel and on an Agilent 2100 Bioanalyzer (Agilent Technologies, Santa Clara, CA, USA).

High-quality RNA samples were sent to Novogen Co. (Beijing, China) for small RNA libraries construction and further sequencing. Briefly, total RNA samples were size fractionated, and small RNAs were used as input for the NEBNext® kit to construct the libraries. Single-end sequencing was performed to produce 50 bp reads (SE50) with 30M reads per sample, using the Illumina NovaSeq6000 platform. RNA-Sequence data were submitted to the NCBI SRA database (accession number PRJNA1498010).

### MicroRNA prediction

Quality control of the reads was performed using FastQC^39^ before and after trimming with TrimGalore^40^ to remove adapter sequences, low-quality reads, and reads shorter than 18 nt. Trimmed reads were collapsed and mapped to the reference genome of *Spodoptera frugiperda (*AGI-APGP_CSIRO_Sfru_2.0, GenBank: GCA_023101765.3) and to Spodoptera frugiperda Multiple Nucleopolyhedrovirus ARG-M genome (GenBank: MW162628.1) using the miRDeep2 mapper module.^41^ Mapping files, trimmed, collapsed reads, and known miRNAs database, including miRbase v22.1 miRNAs and those previously reported by Moné et al.^18^ and by Mahalle et al.^19^, were used to quantify known miRNAs and identify putative novel miRNAs with the quantifier module of MiRDeep2. When the score was equal to or greater than 4, or if the pre-miR had a significant randfold p-value, novel miRNAs were considered valid.

### MicroRNA duplication events identification and analysis

Mature miRNAs with identical sequences were selected and grouped. Multiple sequence alignments (MSA) or pairwise sequence alignments (PSA) of precursors encoding duplicated miRNAs were performed with MAFFT^42^ or Needle,^43^ respectively.

### MicroRNA cluster identification and analysis

MiRNA clusters were identified by looking for two or more pre-miRNAs coded in the same orientation and within a genomic window of 15 kbp. This distance was selected based on known cluster lengths such as miR-1∼miR-133 and miR-2/13/71, which in *S. frugiperda* have a total distance between the starting position of the first miRNA and the end position of the last miRNA of about 15 kpb.

### Differential expression analysis

To assess differential miRNA expression between control and infected conditions, we used the *edgeR* package (v3.40.2) of R (v4.1.0). Raw read counts from miRDeep2 were filtered to retain miRNAs expressed in at least two samples. Data were normalized using the TMM method to account for library size differences. A design matrix modeling the two conditions was used to fit a GLM with estimated dispersion and differential expression tested via quasi-likelihood F-tests. P-values were adjusted using the Benjamini-Hochberg method, and miRNAs with FDR < 0.05 and |log₂FC| > 1 were taken as differentially expressed.

### Target prediction

MiRNA target prediction was performed with three different predictors: miRanda, RNAHybrid and RNA22.^44–46^ Mature differentially expressed miRNA sequences were used as input to predict putative targets in the host 3’UTR transcript sequences and in the SfMNPV Arg-M genome. MiRanda algorithm on Galaxy web server^47^ was used setting a MFE ≤ −20 and Score ≥ 140. RNAhybrid was used locally with default parameters, generating 5000 random sequences and getting a p-value associated with the hybridization, MFE ≤ −20 kcal/mol and p-value < 0.05 was used. RNA22 v2 was used as default, and targets with MFE ≤ −14 kcal/mol were kept. Predictions in the same transcript and in binding positions predicted by two of the three predictors, differing by less than 10 bp were kept. GO term annotations were downloaded from NCBI. KEGG annotation was performed using KAAS. For viral target prediction, CDS and intergenic region sequences were used following the same criteria as before, but using a score ≥ 120 for Miranda as the cut-off. Since there are no transcript sequences of SfMNPV, we used the CDS sequences and the intergenic regions in order to predict putative targets. We used the criteria based on the report of Chen et al., (2013)^48^ that 5’UTR length is 150 nt and 3’UTR length is 400 nt.

For further target characterization, targets were selected based on pathway and GO annotation, keeping those involved in processes previously reported to be relevant in infection. Selected miRNAs and target interactions were plotted using circlize^49^, showing the degree of the selected gene in the PPI as the width of the connector.

### Protein - protein interaction network prediction

The whole proteome of *S. frugiperda* was submitted to STRING^50^ webserver to predict protein - protein interaction (ppi) network, and high-confidence interactions (combined score > 700) were kept. Centrality of each node was calculated using the degree as a criterion, and the top 20% of nodes with the highest degree were considered hub nodes. Targets predicted for each DE miRNA were overlapped with hub nodes to identify targets that were either hubs in the PPI network or direct interactors of those hubs. MiRNA-hub targets, miRNA-non-hub targets that directly interact with hubs and their interactions were calculated and plotted for up-regulated and down-regulated miRNA. Proximity clusters were calculated using the Louvain method, and the global process or pathway was annotated using STRING enrichment calculation based on proteins in each cluster.

The methodology used in this work is shown in Figure 1.

**Figure 1.**
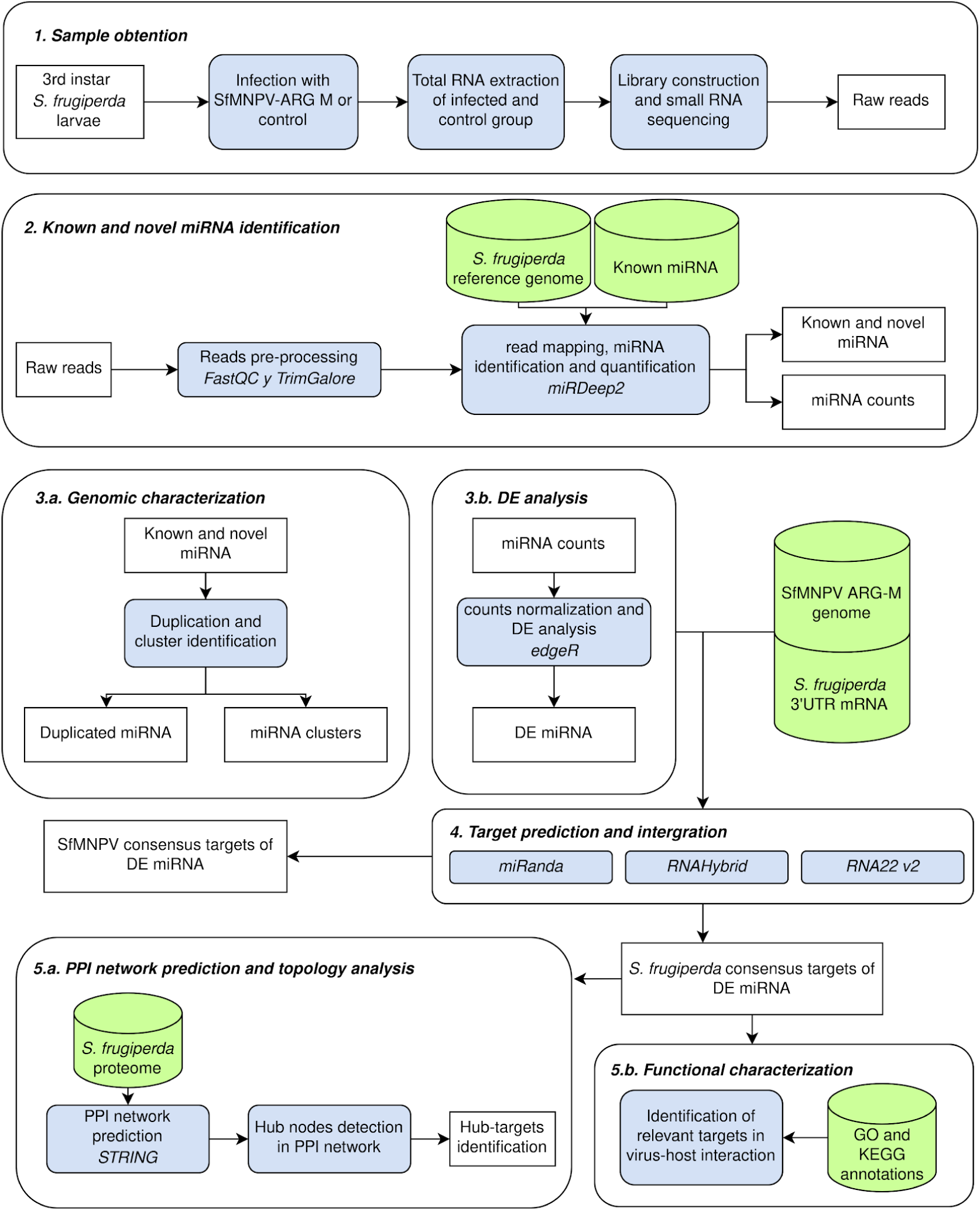
The pipeline followed in this study. Total RNA was extracted from pooled infected and uninfected *S. frugiperda* larvae, followed by small RNA library preparation and Illumina sequencing (1). Trimmed reads were used for known miRNA identification, novel miRNA prediction using the host reference genome, and miRNA quantification (2). Mature miRNA sequences and precursor genomic coordinates were used to identify putative duplication events and genomic miRNA clusters (3a). miRNA expression levels were used for differential expression analysis (3b), and targets of the differentially expressed miRNAs were predicted in both, host and viral mRNAs,, integrating three different methods (4). Functional characterization of host target genes was performed through protein–protein interaction (PPI) network analysis (5a) and Gene Ontology (GO) and KEGG pathway annotations (5b).

## Results

### 1. Small RNA sequencing of *S. frugiperda* samples

SfMNPV-infected and uninfected *Spodoptera frugiperda* larvae were subjected to small RNA-sequencing to identify host-expressed miRNAs. After sequencing, we obtained a total of 155,498,795 reads for the four samples, and after trimming, 112,201,783 trimmed reads were kept. Reads were mapped separately to the host reference genome and to SfMNPV Arg-M isolate genome. Fifty-two percent of the reads mapped to the host genome, while 0.4 % mapped to the virus in the infected samples.

Read length distribution of host-mapped reads was calculated in order to verify that it followed the typical small RNA distribution. The distribution had the characteristic 22 nt peak corresponding to small non-coding RNAs such as miRNAs and siRNAs. We also observed a peak at 28 nt, probably corresponding to piRNA molecules. Read size distribution and specific information of each library is shown in Supplementary Figure S1 and Supplementary Table S1, respectively.

### 2. Known and novel miRNAs were identified in *S. frugiperda* larvae

After verifying the quality of the libraries, trimmed collapsed reads were used as input to identify known and novel miRNA. Since some previously identified miRNAs were not deposited in miRBase v22.1, we built our own *Sfr.* known miRNAs database including those deposited in miRBase, and those reported by Moné et al.^18^ and by Mahalle et al.^19^ This database was used to identify known miRNAs and to distinguish them from putative novel miRNA.

A total of 409 mature miRNAs were found, 360 known miRNAs and 49 novel miRNAs with no homology to any miRNAs deposited in miRBase or previously reported. Among the known miRNA, 247 of them correspond to those deposited in miRBase, 99 to the ones reported by Moné *et al*.^18^ and 14 to the ones reported by Mahalle *et al*.^19^. Detailed information of all detected miRNAs can be found in Supplementary Tables S2 and S3.

We also analyzed the top-expressed miRNAs and compared them to previous reports. We found that Bantam-3p, miR-9c-5p, miR-1a-3p, miR-2766-3p, miR-276-3p, miR-9-5p, miR-10-5p, miR-281-3p, miR-184-3p, miR-263a-5p were among the top 10 most expressed miRNAs (Supplementary Table S4). This result is similar to those previously reported by other authors.^21,23,51^ Novel miRNAs identification and miRNA counts are shown in Supplementary Table S3 and S4.

### 3. Some *S. frugiperda* miRNAs have multiple copies in the genome

MiRNA duplication events in the host genome can help to understand regulatory, and evolutionary dynamics of these regulatory elements.^52^ Duplicated pre-miR can conserve 100% identity in the mature miRNA sequence, or diverge, resulting in paralogs^53,54^. Duplications can be located in the same strand and within the same genomic region, resulting in miRNAs coded in tandem generating a cluster, or can have copies in different chromosomes, after whole-genome or segmental duplication events during evolution.^55–57^ Since this, we wondered whether host miRNAs could have several copies in the genome. We kept predicted novel and known miRNAs whose mature miRNA sequences were identical, but pre-miR coordinates were different. We found 23 duplication events, comprising 59 pre-miRNAs in total. Eighteen duplication events correspond to known miRNA, 9 of them previously reported, and 14 reported in our work. Five duplication events were found among the novel predicted miRNAs. (Figure 2, Supplementary Table S5).

**Figure 2.**
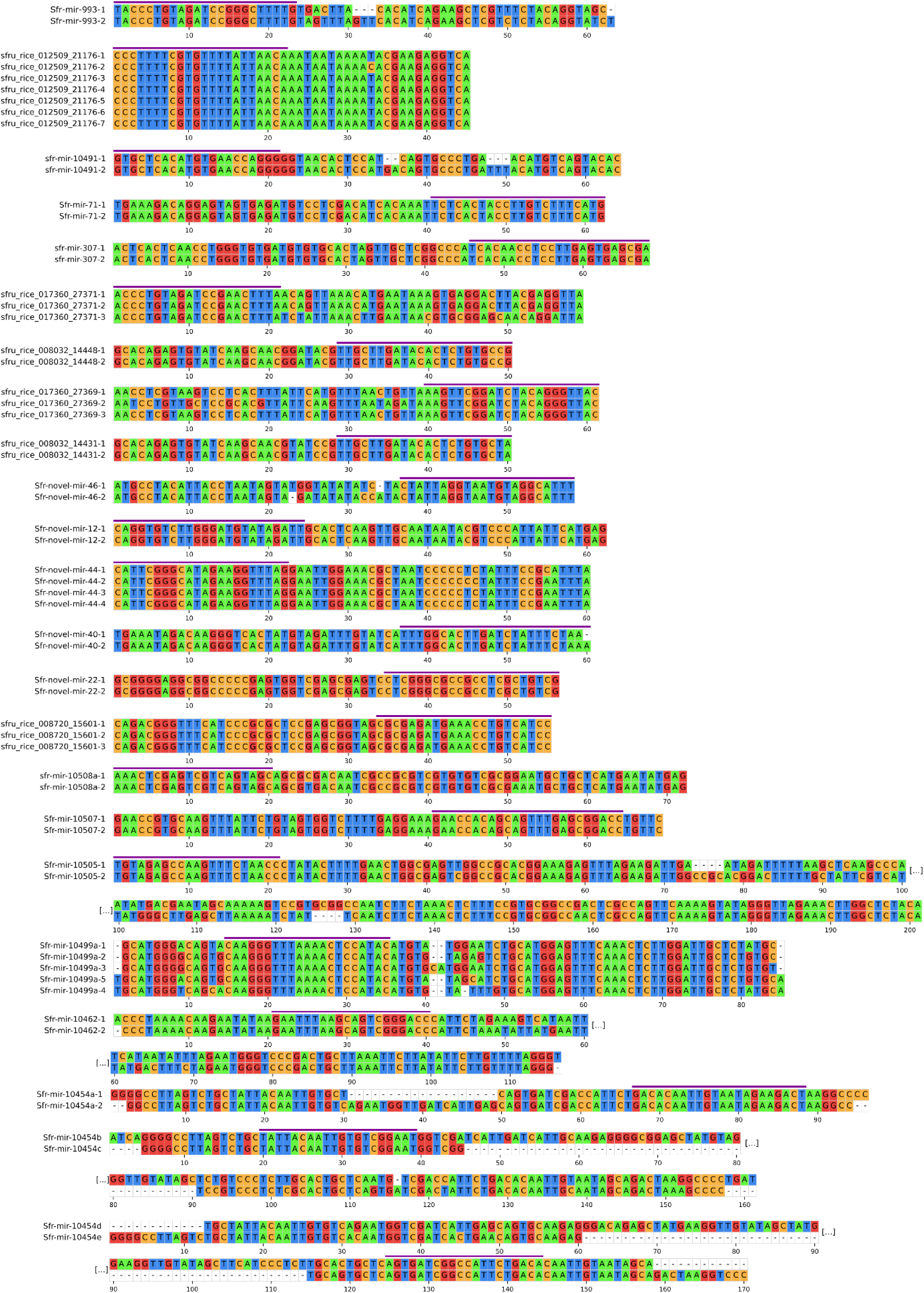
Precursor alignments of duplicated mature miRNA. PSA or MSA of pre-miR encoding the same mature miRNA, mature miRNAs are shown with a purple line.

Regarding known *S. frugiperda* miRNAs miR-71, miR-307, miR-993, they are conserved within other species and duplication events were reported in other organisms,^58–60^ while miR-10454, miR-10462, miR-10499a, miR-10505, miR-10507, miR-10508a are specific from *S. frugiperda* and duplication events were already reported^21^. Other known miRNAs reported by Moné *et al*.^18^ were found duplicated in the genome assembly analyzed in this work. Among them we found duplication events of sfru_rice_012509_21176, sfru_rice_017360_27371, sfru_rice_008032_14448, sfru_rice_017360_27369, sfru_rice_008032_14431, sfru_rice_008720_15601, interestingly finding sfru_rice_012509_21176 coded 7 times in tandem in a genomic region of 4836 bp. We also analyzed sequence conservation between precursors to understand whether both arms or only one arm was conserved. We found some cases with whole precursor duplication, other cases where precursor sequences were not identical but 5p and 3p miRNAs had 100% of identity, and finally, other cases where the miRNAs of only one arm of the precursor had 100% of identity, giving rise to paralogs.

Regarding novel predicted miRNA, we observed that novel-miR-12, novel-miR-22, novel-miR-40, novel-miR-44, and novel-miR-46 are duplicated. Novel-miR-40 has two paralogs, novel-miR-40a and novel-miR-40b, where the sequence is mostly identical, with an extra nucleotide at the 3’ end of novel-miR-40b. Interestingly, we also found novel-miR-44 coded 4 times in tandem, in a genomic region of 1630 bp.

Finally, an interesting finding was that sfru_rice_008032_14448 and sfru_rice_008032_14431 were coded overlapped in opposite strands, with the same situation observed for novel-mir-46-1 and novel-mir-46-2, coding the exact same mature miRNA. This has been previously reported for other miRNAs such as miR-iab-4 and miR-iab-8 in *Drosophila melanogaster.*^61^ In both cases, we found that mature miRNAs generated by pre-miR coded in opposite strands in the same locus shared a high percentage of identity, with 100% identity in the seed sequence.

### 4. *S. frugiperda* miRNAs are coded within miRNA clusters

In addition to duplication events, miRNA clusters contribute to its effect by generating multiple mature miRNAs from a single pri-miRNA transcript.^62^ This can help the host cell to regulate several target genes simultaneously and thereby amplify their regulatory impact.^55,63^ Clusters can be homoclusters, where several pre-miRNAs generate copies of the same mature miRNA, or heteroclusters where pre-miRNAs generate different mature miRNAs. Identifying novel clusters can help to understand the regulatory network of the host. We wondered whether any of the identified miRNAs could be expressed from the same pri-miRNA, so we set out to identify putative clusters. We looked for miRNAs coded in the same strand and chromosome, within a window of 15 kbp. We identified a total of 35 putative miRNA clusters, comprising 115 pre-miR and 141 mature miRNA. The genomic organization of clusters is shown in Figure 3. Detailed information on miRNA clusters can be found in Supplementary Table S6.

**Figure 3.**
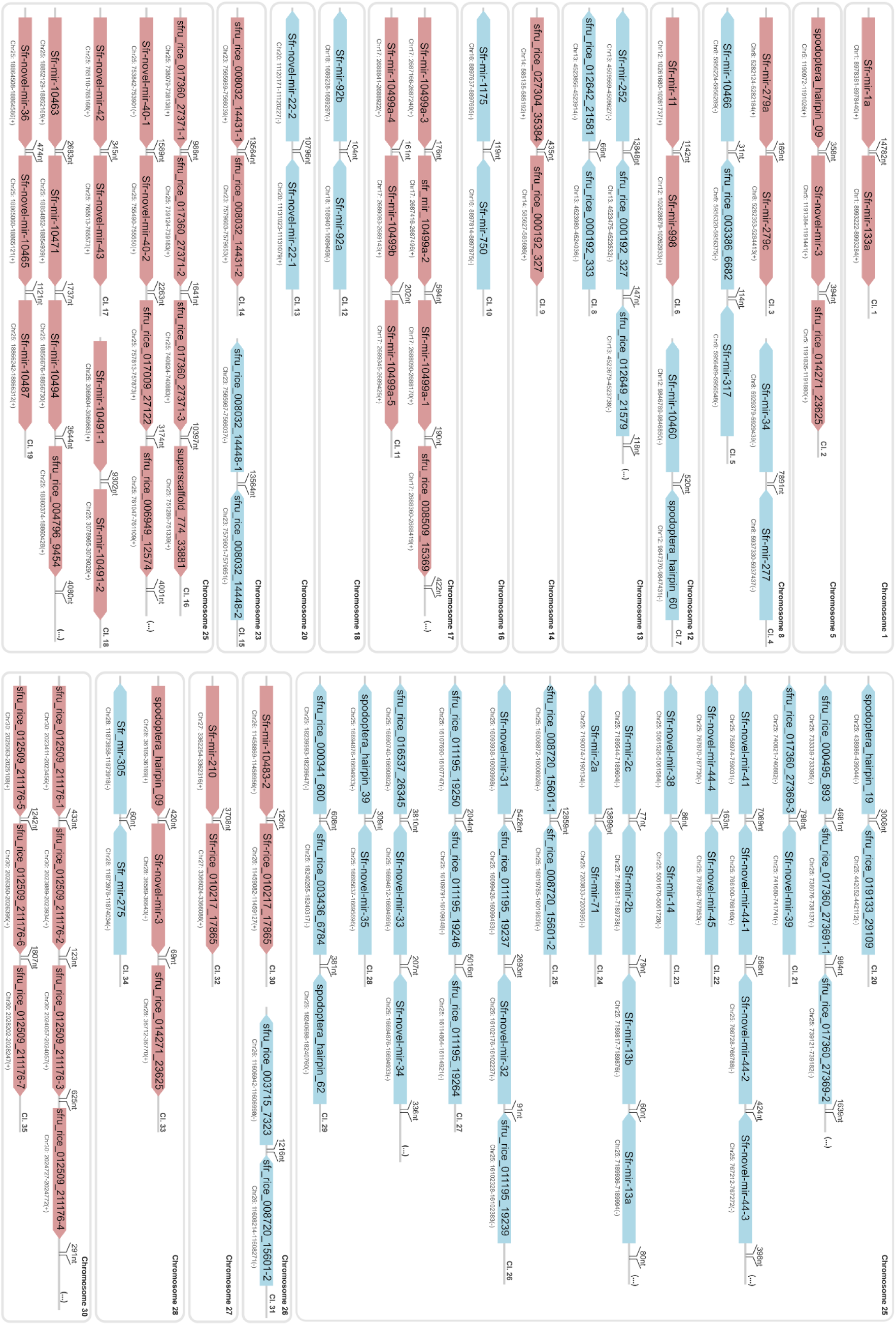
Predicted clusters based on identified miRNAs in *Spodoptera frugiperda*. The figure shows miRNA clusters 1-35 organized by chromosomes. The distance between precursors is shown in each scheme.

We found 9 miRNA clusters that were previously reported in other insect species or metazoa: miR-1a/miR-133a,^64,65^ miR-13∼miR-71,^66^ miR-92 family,^67^ miR-11/miR-998,^68^ miR-279 family,^69^ miR-275/miR-305,^70^ miR-34/miR-277,^71^ miR-79∼miR-9c,^72^ and miR-750/1175.^73^

We also identified 26 putative novel miRNA clusters comprising known and novel miRNAs. Among them, we found that miR-10499b, which belongs to a family with members coded within a cluster according to miRBase annotation, clustered together with sfru_rice_008509_15369 (Cluster 10, Figure 3). We also predicted a putative second copy of mir-10491, generating a putative novel cluster. As previously mentioned, several duplication events that occur in the same strand and in the same genomic region can generate miRNA homoclusters, as is the case of clusters 13, 14, 15, 18, 25 and 35. We detected that some duplication events generated copies in the same strand and genomic region, but also were coded near to another pre-miR, giving rise to heteroclusters. This situation was observed for clusters 16, 17, 21, and 22. Finally, the rest of the predicted clusters consisted of pre-miR that generated different mature miRNAs.

Clusters were found in chromosomes 1, 5, 8, 12, 13, 14, 16, 17, 18, 20, 23, 25, 26, 27, 28 and 30, interestingly finding 15 out of 35 clusters coded in chromosome 25.

A relevant characteristic among miRNA heteroclusters is that some members of the cluster can share sequence similarities, which might reflect evolutionary dynamics and previous duplication events.^67,74,75^ In order to analyze this, we performed multiple sequence alignments (MSA) between mature miRNAs of heteroclusters. MSA of mature miRNAs that have sequence conservation is shown in Figure 4.

**Figure 4.**
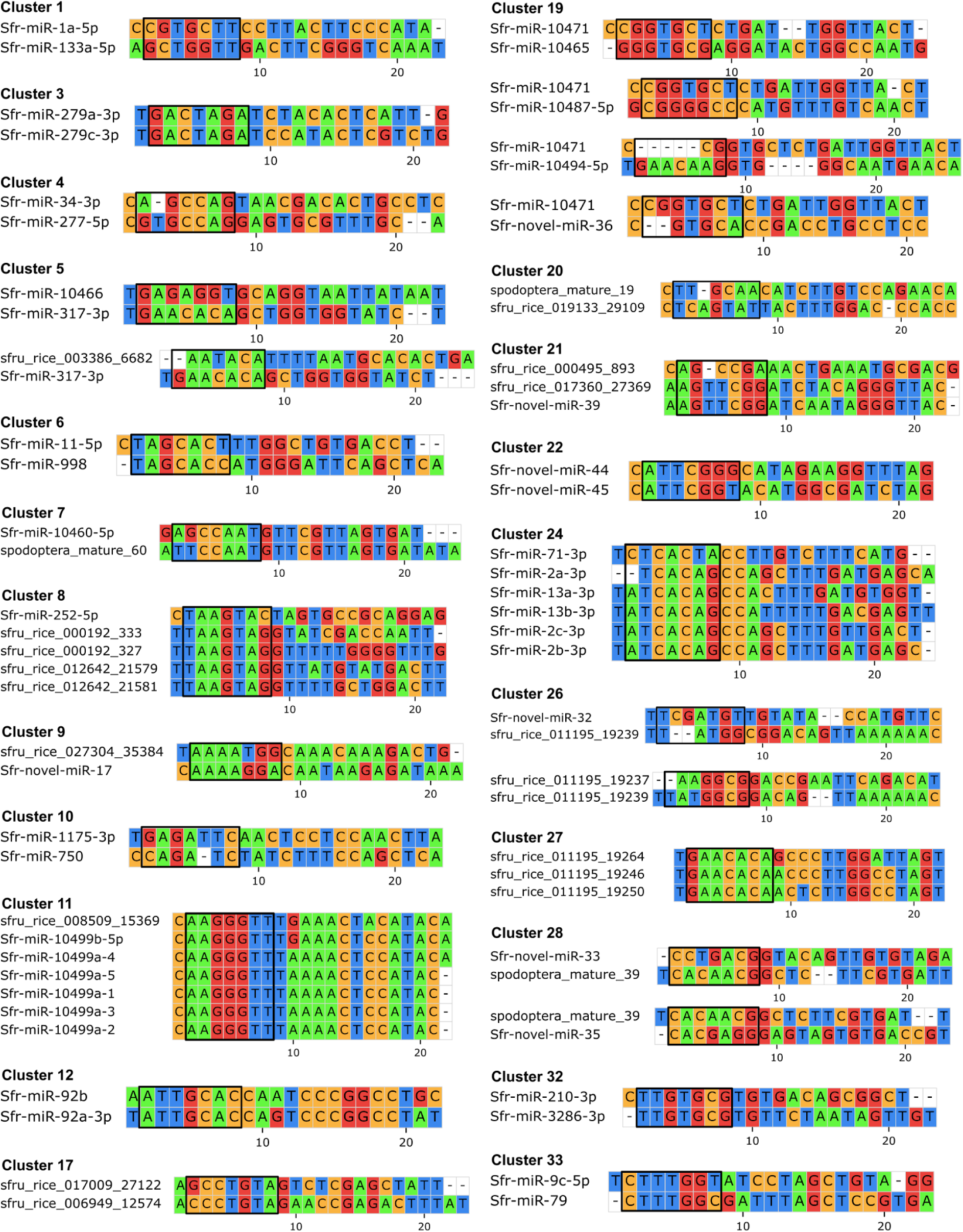
Sequence conservation in miRNA heteroclusters showed sequence conservation between its members. Multiple sequence alignment or pairwise sequence alignments of mature miRNAs of selected clusters. In those cases where sequence was not conserved between all members, but was found in different pairs several PSA are shown for a single cluster.

We found different situations when analyzing sequence conservation, in some cases seed sequence was 100% identical between its members, other cases where seed sequence was similar but no identical, cases where only some members shared seed sequence similarity while the rest did not, seed shifting events, and finally cases where one miRNA shared seed sequence similarity with multiple cluster members, while those members did not share similarity with each other.

For clusters 3, 11, 12, and 27, the seed sequence was 100% identical between its members, suggesting that those miRNAs are part of the same family. This has already been reported for miR-279, miR-92, and miR-10499 families. Moreover, we found that sfru_rice_008509_15369 was coded in a cluster with miR-10499 family, and shared seed sequence, suggesting it might be part of this family too. Also, a novel family consisting of sfru_rice_011195_19264, sfru_rice_011195_19246 and sfru_rice_011195_19250 was identified, where the three members of the cluster 27 shared seed sequence. For clusters 1, 7, 8 and 9 seed sequence similarity was partially found. In cluster 8 also a group of miRNAs shared seed sequence, suggesting that they might be part of the same miRNA family. Analyzing clusters 17, 19, 21, 22 and 24 we found some groups of miRNAs with identical or highly similar seed sequence, but shared no sequence similarity with the rest of the cluster members. For clusters 4, 6, 26, 32, 33 we observed seed shifting events.^76^ In this way, allowing gaps in the alignment, a high percentage of identity could be found between miRNA sequences (Figure 4). We identified several clusters (5, 19, 26, and 28) in which one miRNA shared seed sequence similarity with multiple cluster members, although those members shared no sequence conservation with each other. Pairwise alignment analysis revealed sequence conservation between certain miRNA pairs (Figure 4), while multiple sequence alignment detected no conservation across all members (data not shown). For example, in cluster 5, Sfr-miR-317-3p exhibits sequence conservation with both miR-10466 and sfru_rice_003386_6682, even though these two sequences did not share any detectable similarity. This pattern suggests sequential duplication and divergence, where an intermediate paralog retained similarity to both ancestral and derived copies.^67,75^ Cases where no sequence conservation was found between cluster members were not included in the figure.

Despite our cluster demarcation criteria, we found pairs of neighbouring clusters suspected to be part of a bigger cluster. Clusters 16 and 17 are next to each other in chromosome 25, coded in the same strand, and some of their members share sequence similarities in their seed sequence, suggesting that these two clusters could actually be one cluster. The same happens with members of clusters 21 and 22. And, strikingly, these two bigger clusters were found in the same genomic region in opposite strands. The combined MSA of members of the suggested novel clusters are in Supplementary Figure S3. We also found that although clusters 4 and 5 had been classified as separated clusters using 15 kbp criteria, it has been previously reported in *D. melanogaster* that miR-34, miR-277 and miR-317 are part of the same cluster^77^. A novel finding in this work is that mir-10466 and sfru_rice_003386_6682 might be also part of this cluster (Figure 3).

### 5. MiRNAs expression profile changes in *S. frugiperda* upon SfMNPV infection

To find miRNAs potentially relevant in the infection process we set out to identify those that were differentially expressed (DE) in infected larvae compared to uninfected control. We found 13 DE miRNAs, 4 up-regulated and 9 down-regulated in the infected condition. Up-regulated miRNAs were sfru_rice_007900_14215, Sfr-miR-10483-2-3p, Sfr-novel-miR-44 and sfru_rice_027304_35384, and the down-regulated were sfru_rice_017367_27388, sfru_rice_020251_30137, Sfr-miR-10490-3p, Sfr-miR-10508a-2-3p, sfr-miR-10460-5p, spodoptera_mature_60, sfr-miR-10492b-5p, sfr-miR-10462-5p and Sfr-novel-miR-7 (Figure 5. A, B and C). Raw miRNA counts, differential expression analysis results and PCA of the samples are shown in Supplementary Table S4, Supplementary Table S7 and Supplementary Figure S2, respectively.

**Figure 5.**
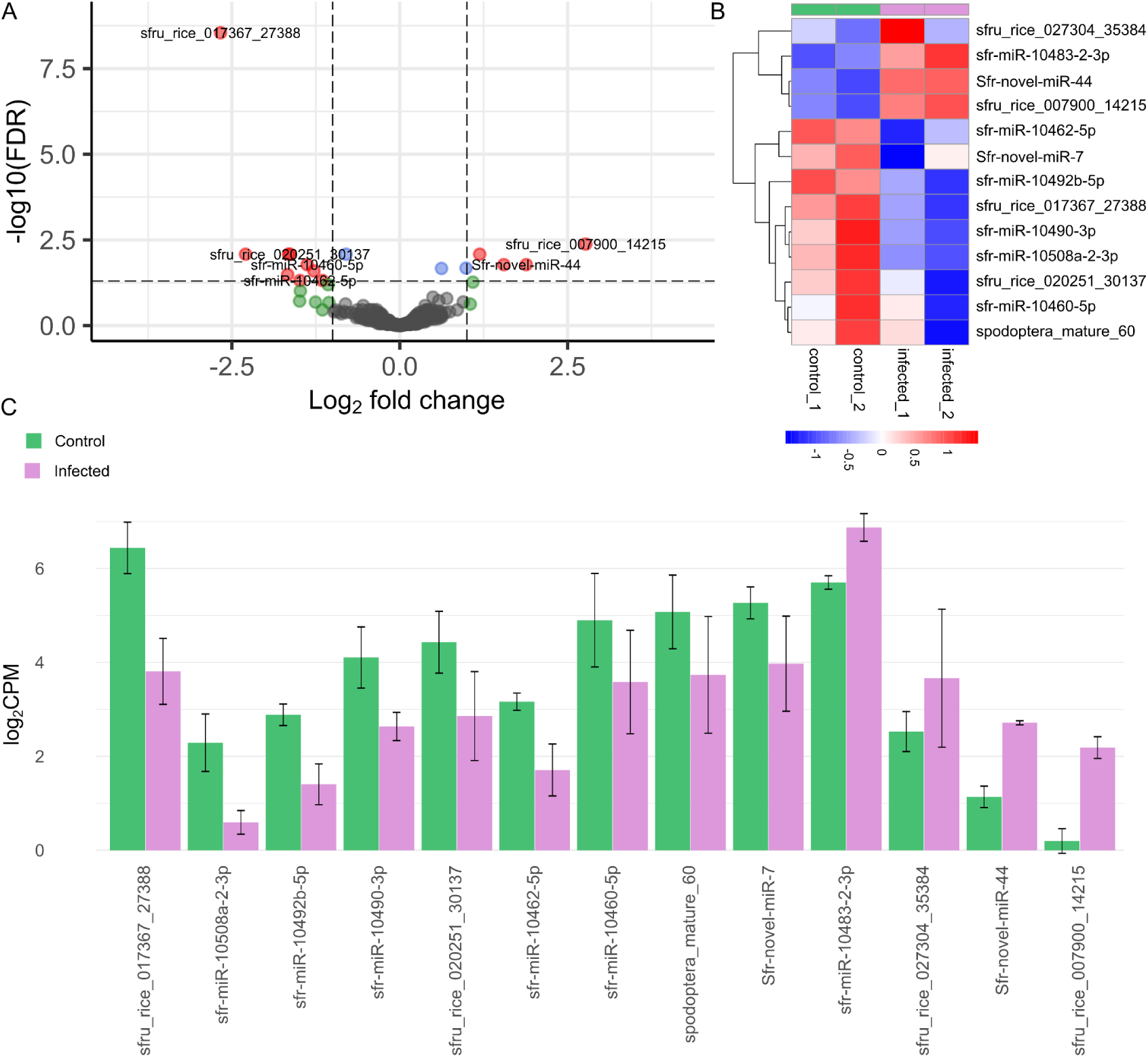
Differential expression analysis of miRNAs. A. Identification of differentially expressed (DE) miRNAs. DE miRNAs are indicated in red. B. Expression of DE miRNAs in different samples. C. Number of reads of DE miRNAs expressed in logCPM in the infected and control group.

Interestingly, among up-regulated miRNA, novel-miR-44 was found to be coded 4 times in the genome, and to be part of cluster 22, while sfru_rice_027304_35384 and miR-10483-2-3p were also generated by pre-miR coded in other miRNA clusters (Supplementary Table S6). When analyzing down-regulated miRNA, we found that both miR-10460-5p and spodoptera_mature_60 were generated by pre-miR members of cluster 7.

### 6. Differentially expressed miRNAs target specific processes and pathways related to virus-host crosstalk

After identifying 13 DE miRNAs, we looked for their putative targets using miRanda, RNA22 V2 and RNAHybrid on *S. frugiperda* 3’UTR of transcripts deposited in RefSeq.

We found 849 targets after keeping binding sites that were predicted by 2 of the 3 predictors, 207 for the up-regulated set, and 542 for the down-regulated miRNAs (Supplementary Table S8). Figure 6. A shows the amount of unique genes targeted by each miRNA, with miR-10492b-5p presenting the highest number of targets and Sfru_rice_020251_30137 with no predicted host targets.

**Figure 6.**
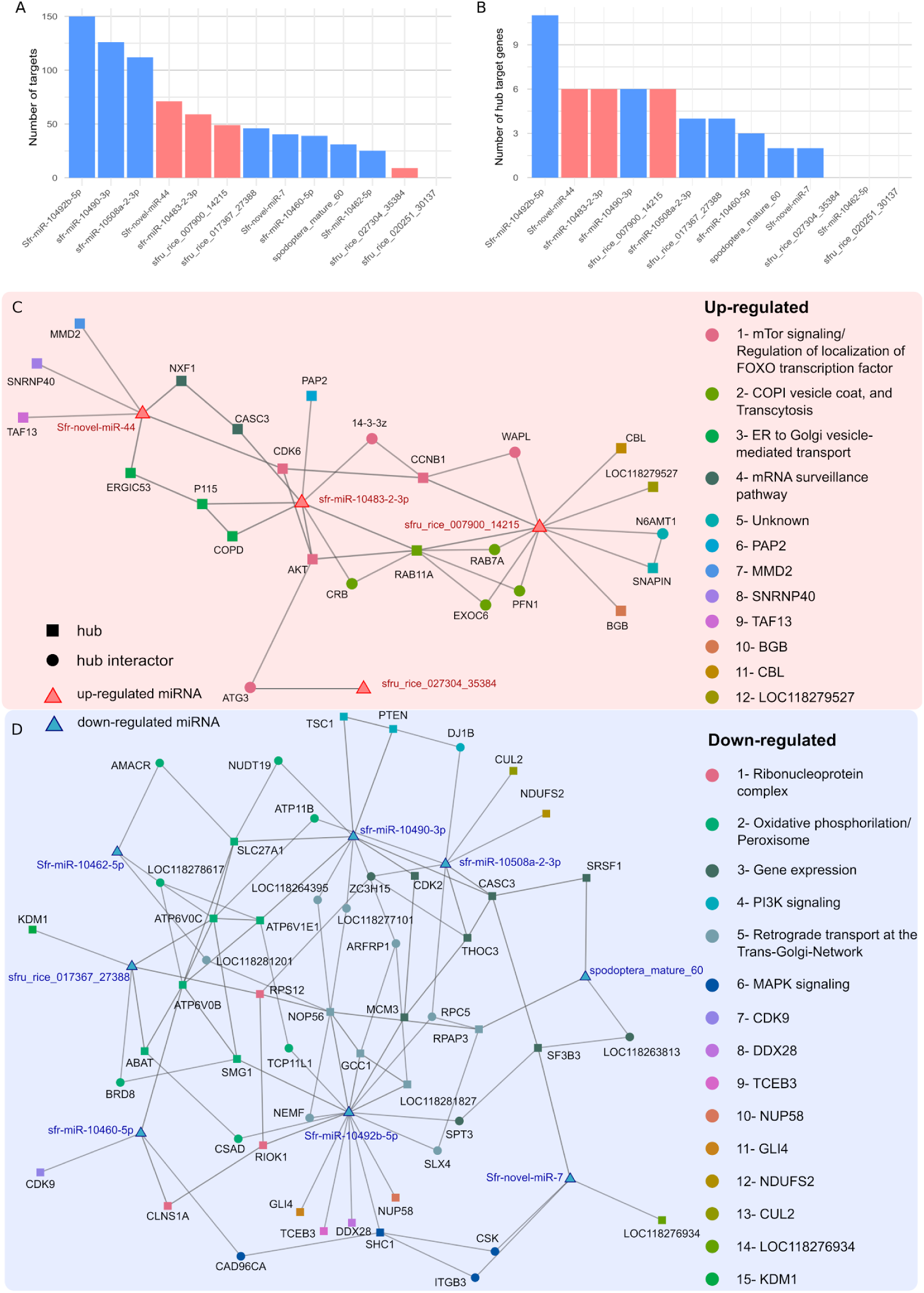
Hub targets in protein - protein interaction network. The figure shows the amount of unique genes (A) and hubs (B) targeted by each miRNA. Up-regulated miRNAs are shown in red, down-regulated miRNAs are shown in blue. C) Up-regulated miRNA - target networks of hub targets, and their interactors that are also targeted by miRNAs from the same group. D) Down-regulated miRNA - target networks of hub targets, and their interactors that are also targeted by miRNAs from the same group. Nodes in each network were clustered, and a general category was assigned.

Several approaches have been used to characterize miRNA targets. Some strategies perform GO or KEGG pathway enrichment analyses to identify overrepresented biological processes. However, these methods do not account for the interdependence among genes.^78^ Alternatively, network based approaches integrate predicted targets into protein - protein interaction (PPI) networks, enabling the analysis of their topological properties. In this context, highly connected nodes (hubs) can be identified, as they are often considered key regulators of biological systems.^79^ In this way, we predicted a PPI network with the whole proteome of *S. frugiperda* using STRING. After keeping high confidence interactions and unique genes to reduce isoforms redundancy we obtained a network with 6620 nodes. We got 1330 hubs as the top 20% of the nodes with the highest degree. Among these, 47 hubs were targets of 10 of 13 DE miRNAs (Figure 6.B). To better understand the interaction between DE miRNAs and their hub targets, we separated them into two groups according to whether they belonged to the up-regulated or down-regulated miRNA groups. Based on the previous PPI network, we analyzed their interaction and included the neighbors of hub nodes that were also targeted by any miRNA from the same DE group in the analysis. These networks are shown in Figure 6.C and D. We found that some hubs were targeted by more than one DE miRNA, such as CASC3 that was predicted as target of down-regulated miR-10490-3p and miR-10508a-2-3p, and RAB11A targeted by up-regulated miR-10483-2-3p and sfru_rice_007900_14215. Interestingly, CASC3 was also targeted by the up-regulated miRNA miR-10483-2-3p. This protein was reported to play a role in non-sense mediated RNA decay and splicing.^80^ Analyzing up-regulated miRNA targets, we found several genes that were reported as relevant to viral infections, or that play a key role in cell biology, therefore playing a role in the infection. Among these targets were Cyclin-B (CCNB1), which regulates G2/M transition;^81^ Cyclin-dependent kinase 6 (CDK6), which facilitates viral DNA replication by interacting with ODV-EC27;^82^ RAC serine/threonine-protein kinase (Akt), which is activated upon baculovirus infection to prevent early cell death,^82^ the transcription initiation factor TFIID subunit 13 (TAF13) which is part of TFIID and has been reported to be exploited by baculovirus to enhance transcription of its own genes by alteration of TBP, the central protein of this complex^83^ and finally, the ras-related protein Rab-11A, which plays a central role in vesicle transport, and has been reported to be exploited by other viruses.^84–86^ When analyzing down-regulated miRNA targets, we also found relevant genes related to cell biology such as cell cycle and transcription regulation. PTEN, a negative regulator of the PI3K/Akt pathway,^87^ the cyclin-dependent kinase 9 (Cdk9) and the transcription elongation factor B polypeptide 3 (TCEB3), which are key regulators of the transcriptional elongation and has been reported to be exploited by other viruses.^88,89^ Cyclin-dependent kinase 2 (Cdk2) that regulates the S phase, reported to be altered upon baculovirus infection^90^ was also a target of this group. Finally, the DNA replication licensing factor (Mcm3), which acts as a helicase to contribute to the DNA replication process,^91^ was also detected as a target.

To identify functional relationships among target genes in both groups, we performed proximity detection using the Louvain method on the corresponding PPI networks. The resulting modules were then compared across networks and visualized by differential node coloring (Figure 6.C and D). We calculated several clusters between targets in each DE group, which were annotated according to their functional enrichment in STRING. Additionally, some singleton clusters were found. For up-regulated miRNA cluster we identified clusters containing genes involved in the mTor signaling pathway and regulation of localization of Foxo transcription factor, as well as clusters related to vesicle transport such as genes involved in COPI coated vesicles and ER to Golgi mediated transport. Finally a cluster linked to the mRNA surveillance pathway was also identified (Figure 6.C).

Among down-regulated miRNA targets, we found additional interesting clusters, two of them related to the regulation of proliferative pathways such as PI3K and MAPK signaling. We also identified a cluster related to gene expression regulation, and another related to Oxidative Phosphorylation and peroxisome, with many members that are subunits of ATPase (Figure 6.D).

Finally, we identified target hubs that did not cluster with any other targets in either group, but are key regulators of many biological processes altered during baculovirus infection. Detailed information of target hubs and the miRNA-target network can be found in Tables S10 and S11.

Even though hub targets allow us to visualize key biological processes regulated by DE miRNA, we wondered whether other targets that might not be hub nodes in the PPI network could be important for the viral infection. To achieve this goal, we used GO and KEGG annotations to classify targets representing biological pathways and processes previously implicated in the regulation of the infection, such as proliferative and apoptotic pathways, DNA damage response (DDR), cell cycle regulation, Toll signaling pathway, JAK/STAT pathway, immune response, 20E and JH signaling pathway, piRNA, siRNA and miRNA pathways (ncRNA), transcriptional pausing, RNA decay, translation regulation, vesicle transport regulation, cytoskeleton remodeling regulation, autophagy, and nucleocytoplasmatic transport regulation. Figure 7 shows the links of the DE miRNAs with their targets associated with these processes. We obtained a total of 110 unique targets involved in these categories, targeted by 12 DE miRNA, 19 of which were also hub nodes in the PPI network.

**Figure 7.**
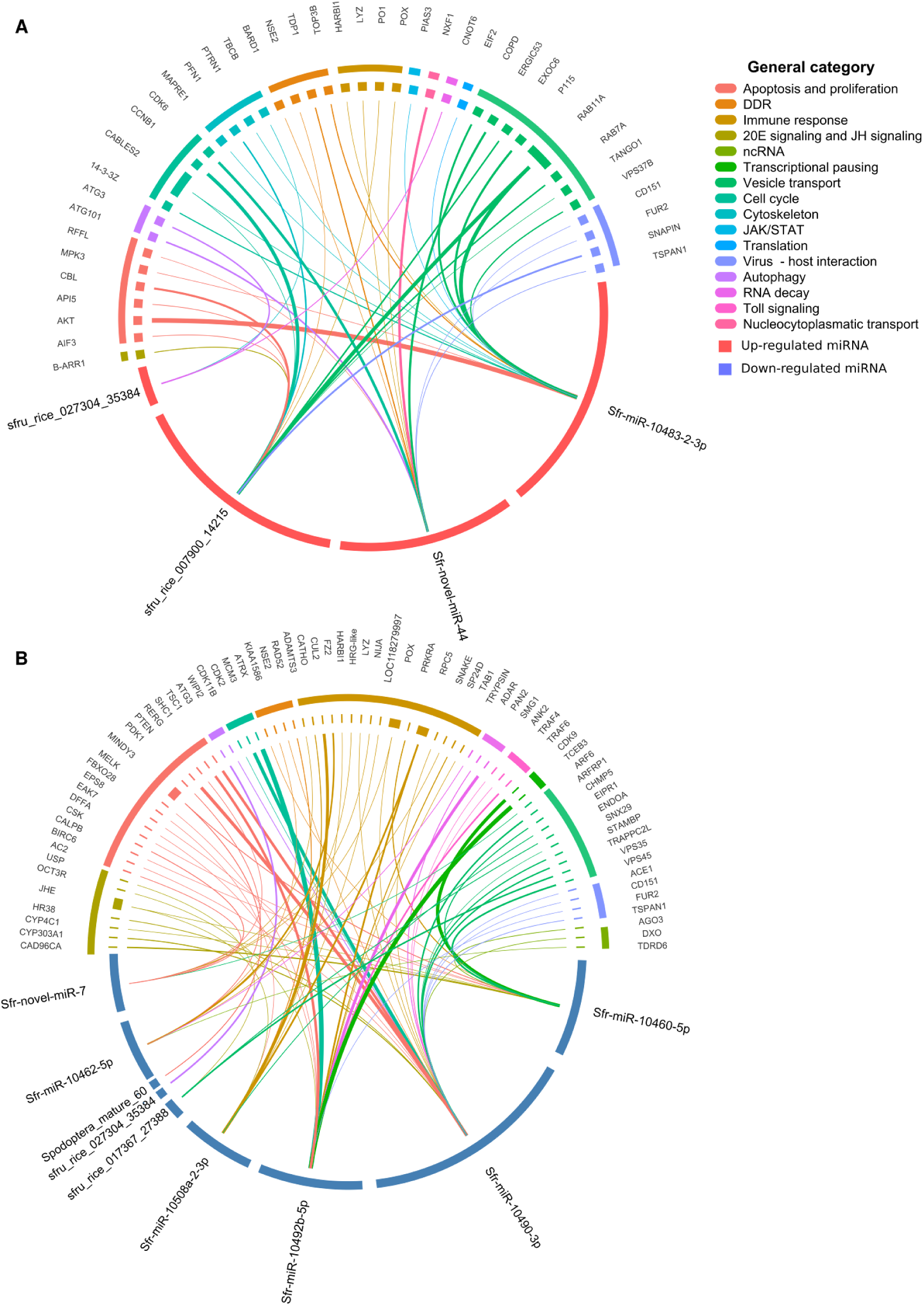
Selected miRNA targets that might be relevant during infection. Circular representation of up-regulated miRNA (A) and down-regulated miRNA (B) and their targets. The general category to which they belong is shown with a different color. The width of the connector indicates the degree of the node in the ppi network, indicating a more central role of those targets with wider connectors.

For the up-regulated miRNA target subset we had 41 unique targets, finding CDK5 and ABL1 enzyme substrate 2 (CABLES2) and RAB11-A targeted by two different up-regulated miRNA. Within this subset of targets, several functional modules were represented. Genes involved in vesicular trafficking were identified, including Rab7A, Rab11A and COPD, central regulators of vesicular trafficking pathways that are widely exploited by viruses. In particular, Rab11A has been extensively demonstrated as a key host factor for viral genome transport and assembly in multiple viruses,^84–86,92^ while Rab7A and COPI components regulate endosomal maturation and Golgi-associated trafficking, respectively, processes known to be critical for baculovirus entry and intracellular transport.^93,94^ Components regulating the balance between apoptosis and proliferation were also detected, notably AKT, a central regulator of the PI3K/AKT signaling pathway.^95^ Cell cycle related targets were present as well, particularly CDK6 and Cyclin B1 (CCNB1), which are involved in the regulation of G1 progression and G2/M transition, respectively.^96^ The JAK/STAT pathway also appears to be modulated through targeting of PIAS3, a known negative regulator of this signaling cascade^9,10,97^. In addition, genes associated with autophagy (ATG3 and ATG101), cytoskeletal organization, and DNA damage response (BARD1, TDP1 and TOP3B) were also identified.^98–100^ Immune related targets were detected, particularly antimicrobial peptides and Phenoloxidase (PO1), a key enzyme in melanization. Finally, several genes potentially involved in virus–host interactions were found, including Furin, TSPAN1, CD151 and SNAPIN, which may contribute to viral entry and infection processes.^101–103^

On the other hand, for the down-regulated miRNA target subset we kept 73 unique targets, finding ubiquitin carboxyl-terminal hydrolase (MINDY3) and DNA-directed RNA polymerase III subunit (RPC5) targeted by two different down-regulated miRNA. MINDY3 is reported to be involved in DDR, apoptosis, and protein quality control,^104^ while RPC5 is a subunit of RNA pol III and might play a role in viral infections.^105^ We found target genes involved in the balance between apoptosis and proliferation, including PTEN,^87^ as well as cell cycle regulators such as CDK2 and MCM3.^96^ The 20E and juvenile hormone axis also appeared to be affected, with two genes that produce JHE targeted by multiple miRNAs, along with USP.^106^ DNA damage response components, including RAD52, ATRX and NSE2, were also identified as targets.^107^ In addition, autophagy-related genes such as WIPI2 were detected. Genes involved in cytoskeletal dynamics and vesicular trafficking were also present, including ARF6, VPS35, VPS45, CHMP5 and SNX29.^108^ Notably, transcriptional regulation was also affected through targeting of CDK9 and TCEB3.^88^ Components of immune signaling pathways were also targeted, including TRAF4 and TRAF6.^109^ Additionally, genes involved in RNA silencing and processing, such as AGO3 and ADAR,^110^ were identified. Finally, several genes potentially involved in virus - host interactions were predicted targets, including FUR2, TSPAN1, ACE1 and CD151.^101–103^ These results show that up-regulated miRNA targets might act over these processes limiting survival, autophagy, and restricting endosomal vesicle trafficking. On the other hand, down-regulated miRNA might derepress targets, generating a complementary effect by limiting PI3K/Akt signaling, controlling cell cycle progression, inducing DDR, enhancing transcription, ncRNA effect and allowing immune signaling. This target subset, including their classification, can be found in Supplementary Table S9.

### 7. Differentially expressed miRNAs target viral genes

Besides host mRNA, DE miRNAs can also target viral mRNAs. This can have a great impact upon infective cycle progression and allow us to better understand viral - host interaction. To study this, we predicted viral targets using the same strategy as with cellular targets. We could identify that up-regulated miRNAs targeted 12 binding sites in the genome, putatively targeting 8 viral mRNAs, and down-regulated miRNAs targeted 31 binding sites in the genome, putatively targeting 25 viral mRNAs. We also found that some down-regulated DE miRNAs (sfru_rice_020251_30137, Sfr-miR-10508a-2-3p, spodoptera_mature_60, sfr-miR-10462-5p) did not have any viral targets, which has been previously reported as possible in other organisms such as the crustacean *M. japonicus* under WSSV infection, where targets of some DE miRNAs were not predicted.^111^ Figure 8.A shows the binding sites of each miRNA in the SfMNPV genome. To further analyze the role of these targets in the viral cycle, we classified them as structural proteins related to the ODV, BV or nucleocapsid, linked to BV or OB production, viral DNA replication and transcription module, host-interaction or unknown function. We found that down-regulated miRNAs targeted genes involved in ODV structure, such as *pif-0, pif-1, pif-3, pif-8, sf58, gp41* and *odv-e66*; in nucleocapsid structure such as *vp1054* targeted in two different binding sites by miR-10490-3p; and in OB structure such as *polh*. Also replication and transcription related genes were predicted as targets, such as *dna helicase* targeted by two different down-regulated DE miRNA*, dna polymerase, lef-3, pkip-1* and *sf53.* Also genes involved with BV production (*sf110)* and OB production (*sf40*) were predicted, and finally genes involved in virus-host interaction such as *gp37* and *p49*. Regarding up-regulated miRNAs we found that the targets were related to the structure of the nucleocapsid such as *pp78/82* and *desmoplakin*, the structure of the ODV such as ODV-EC27 which is also involved in cell cycle regulation by mimicking a cyclin and interfering with G1/S transition, replication and transcription module such as *lef-1* and *lef-9* and virus-host interaction genes such as *chitinase* and *p13.* Viral targets of DE miRNAs prediction can be found in Supplementary Table S12.

**Figure 8.**
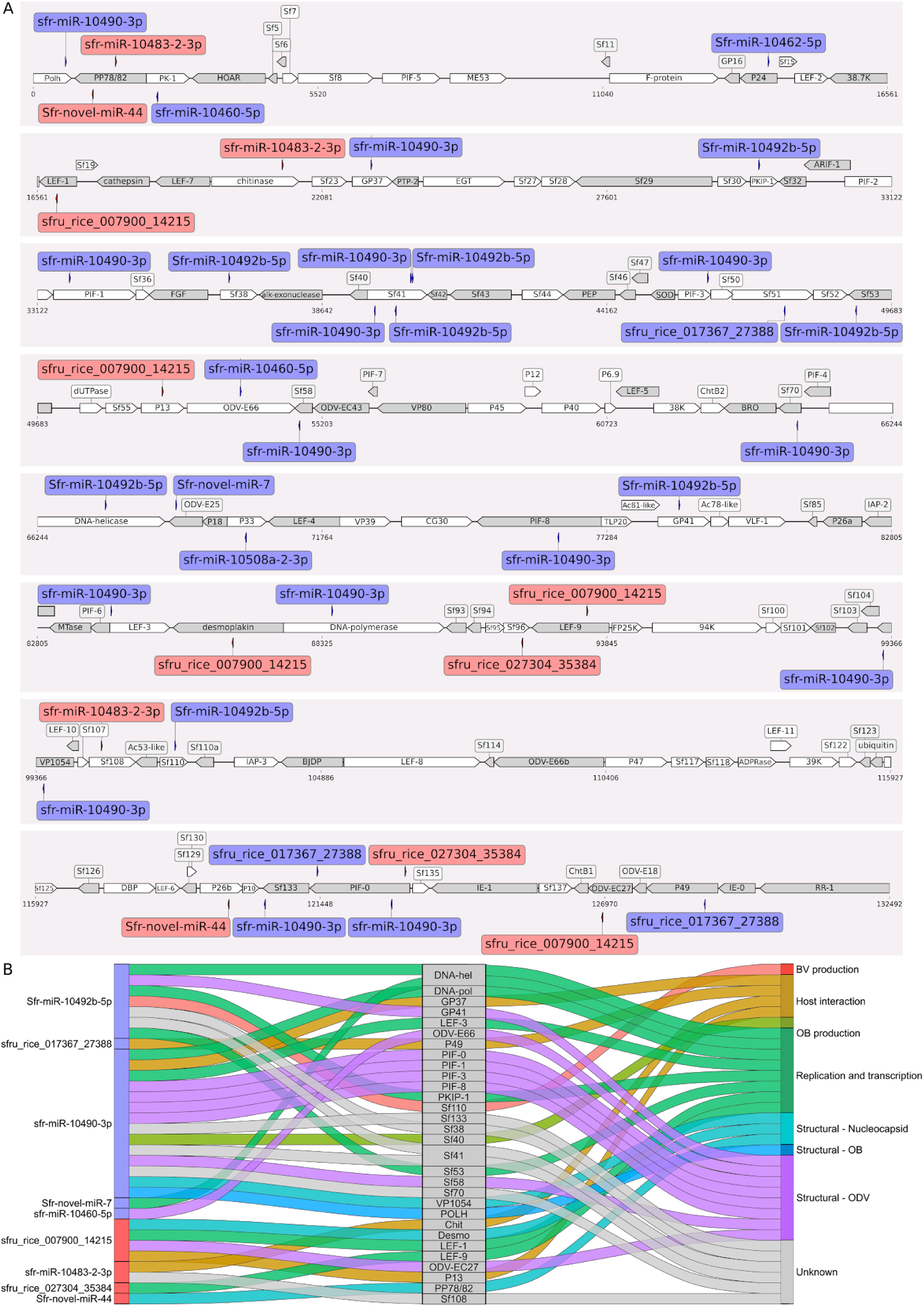
Viral targets of host DE miRNAs. (A) Distribution of DE miRNAs binding sites across the viral genome. Sites on the direct and complementary strands are shown above and below the genome line, respectively. miRNAs are colored by expression (red, up-regulated; blue, down-regulated). (B) Sankey diagram of miRNA–target–functional category associations. Functional categories were assigned from annotation and literature; connectors are colored by target category.

## Discussion

Several papers have previously described changes in microRNA expression in *S. frugiperda* upon baculovirus infection^23–25^ and other experimental conditions,^18–22^ highlighting the miRNA-mediated post-transcriptional regulation in insect biology and host - pathogen interaction. In this study, we reported for the first time a comprehensive characterization of the *S. frugiperda* miRNAome from whole larvae after infection with SfMNPV, including the analysis of duplication events and clustered miRNAs. In addition, we identified differentially expressed miRNAs during infection with the Argentinean isolate SfMNPV-Arg M and explored their potential biological relevance through target prediction, protein–protein interaction network analysis, and viral target prediction. Together, these analyses provide an expanded overview of the *S. frugiperda* miRNA repertoire and identify candidate miRNAs potentially associated with baculovirus infection.

Our genomic analysis revealed the presence of 23 duplication events comprising 59 pre-miRNAs and a total of 115 pre-miRNAs distributed across 35 clusters, including 5 homoclusters and 30 heteroclusters, four of them containing multiple copies of specific miRNA members within the *S. frugiperda* miRNAome. Several miRNAs previously reported by Moné et al.^18^ were also identified as duplicates. Although these duplication events were not detected by the authors, differences might be associated with the differences in genome assemblies used in both studies. Duplicated and clustered miRNAs displayed different levels of sequence conservation, with some copies retaining conservation over the whole mature miRNA sequence, whereas others conserved only the seed region. Within homoclusters, not only mature miRNAs but also pre-miRNA sequences showed high levels of identity, as observed for sfru_rice_012509_21176, which was present in seven tandem copies forming cluster 35. Also, members of clusters 11 and 27 exhibited strong conservation across the complete mature miRNA sequence, whereas cluster 8 showed conservation mainly restricted to the seed region. In clusters 21 and 22, only some members retained highly conserved seed sequences while others diverged, and cluster 28 showed limited sequence conservation among its members. In addition, we identified sense/antisense arrangements, including the duplicated novel-miR-46-1 and novel-miR-46-2 loci resembling the conserved iab-4/iab-8 miRNA locus previously described in insects.^61^ Interestingly, homoclusters 14 and 15, each containing two copies of the same miRNA, were also organized in sense/antisense orientation, further highlighting the complexity of miRNA genomic organization in this species. These genomic arrangements may suggest additional layers of regulatory complexity, since duplicated miRNAs could contribute to increased miRNA dosage, while antisense miRNAs have been proposed to regulate sense miRNA expression or act on common targets.^61,112^ Similar duplication events^58–60^ and genomic arrangements^64–73^ have been described in other species and are thought to reflect the dynamic evolution and putative functional adaptation^55,63^ of miRNA families through local duplication events and subsequent diversification.^52–57^ In particular, clustered miRNAs may arise through tandem duplication or genomic rearrangements, contributing to the expansion and structural organization of the miRNAome in lepidopteran species. We identified 13 differentially expressed miRNAs in response to SfMNPV-Arg M infection, suggesting that baculovirus infection induces substantial changes in the post-transcriptional regulatory landscape of *S. frugiperda* larvae. Similar modulation of insect miRNAs following baculovirus infection has been previously reported in *S. frugiperda* cells and other lepidopteran hosts, supporting the involvement of miRNA mediated pathways during host - virus interactions.^23–36^ Some of the differentially expressed miRNAs identified in this study, such as sfru_rice_017367_27388, Spodoptera_mature_60, miR-10460-5p and miR-10483-2-3p have also been reported to be differentially expressed in other physiological or experimental conditions in insects,^18,19^ suggesting that these molecules may participate in broader regulatory processes beyond baculovirus infection. Although the specific biological roles of many insect miRNAs remain poorly characterized, the differential expression observed in this study suggests that these molecules may participate in the regulation of cellular processes associated with viral infection and host response. Interestingly, some differentially expressed miRNAs were also located within clustered genomic regions. For example, the down-regulated miR-10460-5p and Spodoptera_mature_60 belong to the same cluster and were also reported as co-differentially expressed in a previous study,^19^ whereas up-regulated miR-10483-2-3p, sfru_rice_027304_35384, and novel-miR-44 were associated with clustered arrangements, particularly in the case of novel-miR-44, which was present in multiple miRNA copies. However, the functional implications of this genomic organization remain unclear. Interestingly, in some clusters only a subset of members was differentially expressed. Similar observations have been previously reported and may reflect differences in the processing efficiencies of precursors coded within the same miRNA cluster, suggesting a post-transcriptional regulation. ^113,114^ In our analysis, this pattern was only observed among up-regulated DE miRNAs. Baculoviruses are known to use host machinery to express their own miRNA,^115^ and BmNPV has been reported to alter host miRNA biogenesis through the regulation of the Exportin-5 cofactor Ran.^116^ In addition, several viruses have been shown to modulate miRNA biogenesis by affecting proteins involved in miRNA processing during infection.^117–119^ Taken together, these observations open the possibility that modulation of host miRNA biogenesis machinery could contribute to the differential expression patterns observed among members of the same miRNA clusters during infection. Based on these observations, we further analyzed the predicted host and viral targets of the differentially expressed miRNAs to better understand their potential roles during infection.

Functional characterization of the predicted targets of the DE miRNAs revealed several genes and pathways potentially associated with baculovirus infection and host response. Given the limitations of GO and KEGG enrichment analyses in non-model organisms due to incomplete annotation and the large number of predicted targets, protein–protein interaction (PPI) network analysis was used as a systems-level approach to identify functionally connected genes and potential regulatory hubs within the predicted target set.^78^ This analysis allowed us to identify highly connected nodes (hubs),^79^ related to processes previously implicated in baculovirus infection, including transcriptional regulation, RNA processing, intracellular trafficking, cell cycle regulation, apoptosis, and signaling pathways. Moreover, several targets clustered within interconnected subnetworks, suggesting the presence of functionally related modules potentially affected during infection. The convergence of several predicted targets in pathways related to these processes supports that miRNA-mediated regulation may contribute to the complex balance between host antiviral responses and baculoviral replication strategies.

Several predicted targets identified in our analysis are involved in pathways known to be manipulated during baculovirus infection. Up-regulated miRNAs were predicted to target genes associated with the PI3K/Akt signaling pathway, including AKT, a central regulator of proliferation and survival previously implicated in baculovirus infection.^10^ Since baculoviruses commonly exploit proliferative pathways to support viral replication and dissemination,^82,120,121^ repression of these targets by up-regulated miRNAs could represent a potential antiviral response limiting cell survival and proliferation. In contrast, down-regulated miRNAs were predicted to target PTEN, a gene that has been reported to play an antiviral role during baculoviral infection.^10,122,123^ Derepression of PTEN expression during infection could further contribute to the inhibition of proliferative signaling, suggesting that both up- and down-regulated miRNAs may converge toward modulation of the same pathway through regulation of distinct components with complementary effects. A similar pattern was observed for cell cycle regulation. Up-regulated miRNAs targeted CDK6 and CCNB1, genes involved in G1/S and G2/M progression, respectively; whereas down-regulated miRNAs targeted CDK2 and MCM3, also associated with DNA replication and cell cycle progression. Together, these findings suggest a coordinated miRNA-mediated regulation of multiple stages of the cell cycle during infection. In particular, CDK6 has been reported to interact with the viral cyclin ODV-EC27, an interaction associated with host DNA synthesis shutdown and viral replication.^82^ Additional targets associated with DNA damage response pathways, including BARD1, TDP1, RAD52, ATRX, and NSE2, were also identified among up-regulated and down-regulated miRNA targets, suggesting that both repression and derepression of DDR-related genes may occur simultaneously during infection. Since baculoviruses are known to manipulate DDR signaling to promote replication and regulate apoptosis and cell cycle progression,^124–126^ these interactions may reflect a dynamic balance between host antiviral defense mechanisms and viral exploitation of host replication machinery.

Autophagy, vesicle trafficking, and intracellular transport were also among the processes potentially regulated by differentially expressed miRNA. Several predicted targets, including RAB7A, RAB11A, ARF6, VPS35, VPS45, CHMP5, SNX29, ATG3, ATG101, and WIPI2, have previously been associated with baculovirus infection.^10,93,127–133^ Interestingly, both up and down-regulated miRNAs converged on these pathways while targeting distinct molecular components. Up-regulated miRNAs targeted genes such as RAB11A which has been reported to play a role in baculovirus entry,^127^ ATG3 and ATG101 which has been reported to mediate BmNPV infection,^128,129^ potentially restricting vesicular trafficking and autophagy-related processes exploited during baculovirus infection. Conversely, down-regulated miRNAs targeted genes including VPS35, VPS45, ARF6, and WIPI2, whose derepression could maintain or enhance trafficking and autophagy functions during infection. Rather than indicating strictly antagonistic effects, this pattern may reflect fine regulatory adjustments affecting different stages or branches of these pathways. Since autophagy and vesicle trafficking are known to exert both antiviral and proviral functions during baculovirus infection,^10,93,127–133^ simultaneous repression and derepression of distinct pathway components could contribute to a tightly regulated balance between viral replication requirements and host defense responses.

A similar convergence was observed in immune and regulatory pathways. Up-regulated miRNA targeted genes associated with melanization and immune defense, including phenoloxidase-1 (PO1) and antimicrobial peptides, potentially contributing to suppression of specific immune responses during infection. Also PIAS3, a negative regulator of the JAK/STAT signaling pathway, was targeted. Since this pathway has been reported to play an immune role,^9,10,97^ up-regulated miRNA targeting its negative regulator could imply the activation of the pathway. In contrast, down-regulated miRNA targeted immune signaling regulators such as TRAF4 and TRAF6, whose derepression could enhance immune-related signaling pathways. Likewise, transcriptional and RNA-processing regulators including CDK9, AGO3, and ADAR were specifically associated with down-regulated miRNA, suggesting that derepression of these factors may favor transcriptional activity and RNA-mediated regulatory responses during infection. Interestingly, AGO3 has previously been reported as up-regulated during baculovirus infection,^110^ while antiviral roles have also been described in other lepidopteran viral systems,^134^ supporting the possible involvement of RNA silencing pathways during SfMNPV infection. Hormonal regulation pathways also appeared to be affected by down-regulated miRNA, particularly through targeting of USP, a key transcriptional regulator of 20E signaling, and JHE genes involved in degradation of active JH form. JH and 20E axes are known to modulate antiviral responses, apoptosis, and cell proliferation during infection,^11,12,34,106^ promoting a 20E-immunestimulatory landscape. At the same time, JH generates an antagonistic effect and is also altered upon baculoviral infection by the *egt* gene by inactivating 20E action. These results suggest that miRNA-mediated regulation may contribute to broader physiological and endocrine reprogramming during SfMNPV infection, possibly countering the viral *egt* gene effect and generating a putative antiviral effect.

Finally, several genes potentially involved in virus - host interactions were identified among the predicted targets, including the furin-like protease, FUR2, TSPAN1, CD151 and SNAPIN, all of which have been associated with viral entry, trafficking, or infection processes in different viral systems.^101–103,135^ Together, these findings suggest that differentially expressed miRNA may participate in a complex and multilayered regulatory network during SfMNPV infection, in which both repression and derepression of distinct pathway components contribute to the modulation of apoptosis, proliferation, autophagy, intracellular trafficking, immune signaling, hormonal regulation, and RNA-mediated regulatory mechanisms.

In addition to host targets, several differentially expressed miRNA were predicted to target viral genes associated with viral entry, trafficking, nucleocapsid transport, replication, transcription, and ODV/OB assembly, suggesting a potential direct contribution of host miRNAs to the regulation of the viral cycle. Interestingly, some miRNA simultaneously targeted host and viral genes involved in related biological processes, raising the possibility of coordinated regulation across host–virus interaction pathways. Among the predicted viral targets, we identified several genes directly involved in host manipulation and viral infection processes, including P49, an apoptosis suppressor previously implicated in baculovirus infection.^136^ Down-regulated miRNAs were predicted to target several per os infectivity factors (PIFs), as well as late-expressed genes encoding ODV structural proteins, nucleocapsid-associated proteins such as VP1054, and very late genes including POLH. In addition, genes involved in replication, transcription, and early stages of infection were also identified among the predicted targets. Up-regulated miRNAs were likewise predicted to target late and very late viral genes associated with nucleocapsid structure and transport, including ODV-EC27, an early viral gene involved in cell cycle regulation, as well as genes belonging to the viral replication and transcription machinery. Interestingly, ODV-EC27, also reported as a viral cyclin with a similar function to CCNB1, was targeted by the same miRNA as its cellular counterpart (sfru_rice_007900_14215), and is known to interact with CDK6, also a target of up-regulated miRNA. This interaction was reported to be associated with the shutdown of host DNA synthesis and the initiation of viral replication.^82^ Therefore, targeting these genes by up-regulated miRNA could prevent this process and generate a putative antiviral effect. Interestingly, two late genes involved in host manipulation, CHITINASE and P13, were also identified as predicted targets. Together, these findings suggest that host miRNAs could potentially affect multiple stages of the baculoviral cycle, including viral entry, replication, intracellular trafficking, structural assembly, and modulation of host physiology.

Although the analysis presented here is mainly based on bioinformatic prediction and the putative biological roles of the DE miRNA should be interpreted with caution, the results obtained are promising and provide novel evidence derived from whole *S. frugiperda* larvae infected with an Argentine SfMNPV isolate. Moreover, this work provides an initial framework for understanding the possible regulatory role of these miRNAs during viral infection. Further experimental studies will be necessary to validate both the predicted miRNA - mRNA interactions and the biological relevance of these miRNAs during SfMNPV infection in *S. frugiperda*. Overall, these findings expand the current knowledge of the *S. frugiperda* miRNA repertoire and provide a basis for future studies on miRNA regulation during baculovirus infection.

## Data Availability

The data sets generated in this study are deposited in the National Center for Biotechnology Information (NCBI) repository under accession no. PRJNA1498010

## Supplementary data

Read size distribution, PCA analysis of samples, MSA of combined miRNA clusters 16 and 17, and 21 and 22 are available in Supplementary figures (PDF). All miRNA detected, novel miRNA predicted, raw miRNA counts, duplicated miRNA detected, miRNA clusters prediction, differential expression analysis results, host target prediction, selected relevant targets, hub targets, miRNA-target network and viral targets prediction are available as Supplementary tables (xls).

## Acknowledgment

We acknowledge Instituto de Microbiología y Zoología Agrícola (IMyZA), Centro de Investigaciones en Ciencias Agronómicas y Veterinarias (CICVyA), Instituto Nacional de Tecnología Agropecuaria (INTA), Buenos Aires, Argentina for the provision of *Spodoptera frugiperda* larvae.

## Author Contributions

Gómez Bergna, SM participated in the experimental design, performed experiments, analyzed data and wrote the manuscript. Amorós Morales, LC; Gonzalez Abad, A; Vilches, J analyzed data and helped with data visualization; Tongiani, SE and Salvador, R helped with insect infection assays and helped with insect rearing; Pidre, ML; Romanoski, V contributed to the conception of the study; Ferrelli, ML participated in study conception, experimental design and supervised the study.

## Funding sources

This research was supported by grants from Foncyt (Agencia Nacional de Promoción de la Investigación, el Desarrollo Tecnológico y la Innovación, PICT 2020-2466 and PICT 2021-00969) and CONICET (PIP 2789) to MLF.

## Notes

### Competing Interest Statement

The authors have declared no competing interest.

