## Supplementary Figures for "MicroRNAome of *Spodoptera frugiperda* in Response to SfMNPV Infection"

**Figure S1. Read length distribution of reads mapped to hsot genome.**

**Figure S2. PCA of miRNA expression in samples.**

**Figure S3. MSA of mature miRNA of clusters 16 and 17, and of clusters 21 and 22.**

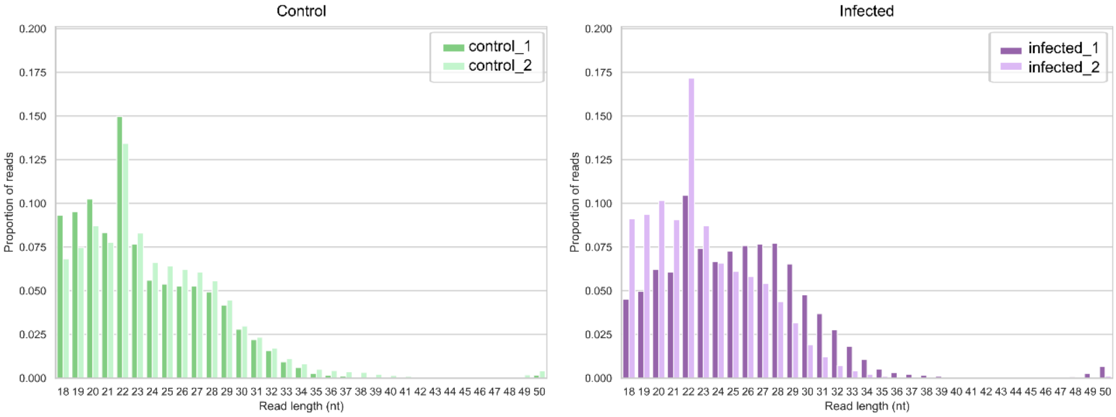


**Figure S1.** Read length distribution of reads mapped to host genome in control simples and infected simples.

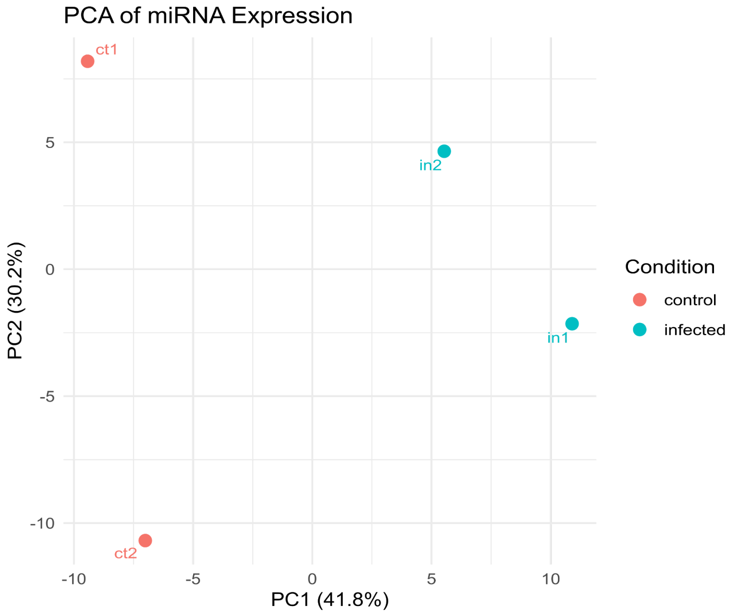


**Figure S2.** PCA of miRNA expression in samples.

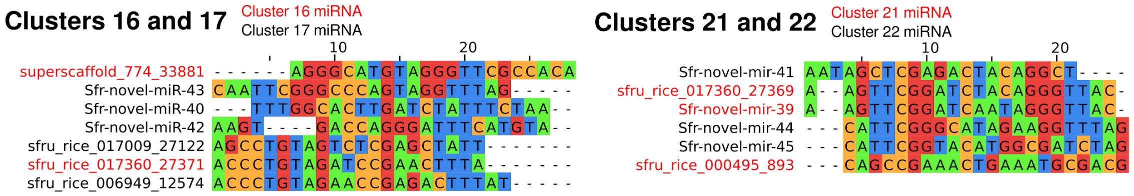


**Figure S3.** Mature miRNA alignments of members of clusters 16 and 17, and clusters 21 and 22.
